# SoyOD: An Integrated Soybean Multi-omics Database for Mining Genes and Biological Research

**DOI:** 10.1101/2024.09.19.613982

**Authors:** Jie Li, Qingyang Ni, Guangqi He, Jiale Huang, Haoyu Chao, Sida Li, Ming Chen, Guoyu Hu, James Whelan, Huixia Shou

**Author notes:** Equal contribution (Jie Li, Qingyang Ni). Correspondence authors. (Shou H), (Hu G), (Chen M).

## Abstract

Soybean is a globally important crop for food, feed, oil, and nitrogen fixation. A variety of multi-omics research has been carried out generating datasets ranging from genotype to phenotype. To utilise this data, a soybean multi-omics database that has broad data coverage and comprehensive data analysis tools would be of value for basic and applied research. We present the soybean omics database (SoyOD), which integrates significant new datasets with existing public datasets for the most comprehensive collection of soybean multi-omics information. Compared to the existing soybean database, SoyOD incorporates an extensive collection of novel data derived from the deep-sequencing of 984 germplasms, 162 novel transcriptome datasets from seeds at different developmental stages, 53 phenotypic datasets, and over 2500 phenotypic images. In addition, SoyOD integrates existing data resources, including 59 assembled genomes, genetic variation data from 3904 soybean accessions, 225 sets of phenotypic data, and 1097 transcriptomic sequences covering 507 different tissues and treatment conditions. SoyOD is a novel tool, as it can be used to mine and analyze candidate genes for important agronomic traits, as shown in a case study on plant height. Additionally, powerful analytical and easy-to-use toolkits enable users to easily access the available multi-omics datasets, and to rapidly search genotypic and phenotypic data in a particular germplasm. The novelty, comprehensiveness, and user-friendly features of SoyOD make it a valuable resource for soybean molecular breeding and biological research. SoyOD is publicly accessible at https://bis.zju.edu.cn/soyod.

## Introduction

Soybean (*Glycine max* (L.) Merr.) is an annual herbaceous crop in the legume family. It is an important source of plant-derived oil, protein and biologically fixed nitrogen. Soybean originated in China and has been cultivated for more than 5000 years. Cultivated soybean is domesticated from wild soybean (*Glycine soja* Sieb. and Zucc.) and is now a global agronomic crop [1]. Over time, cultivated varieties with a range of desired agronomic traits have been developed to suit human needs. However, there are increasing global demands for soybeans, with China alone importing more than 80 million tons annually for food, feed, and industrial uses. These demands underscore the need for developing new soybean varieties with improved yield, quality, and stress tolerance to unfavourable and changing environmental conditions. Since the release of the first soybean reference genome from the variety Williams 82 in 2010 [2], significant progress has been made in soybean genomics research. To date, researchers have successfully assembled the chromosomal-level genomes of more than 50 diverse soybean varieties. [3, 4]. Several complete gap-free genomes (telomere-to-telomere, T2T) have been published [5–9]. Accompanying this are extensive datasets on the genetic variation from thousands of soybean accessions and other omics datasets, including genome resequencing, pan-genomes, transcriptomes, phenomes and more. These resources have greatly accelerated soybean functional genomics research and molecular breeding [10].

To integrate multi-omics data, several soybean multi-omics databases have been established, such as Soybase [11], SoyKB [12], SoyOmics [13] and SoyMD [14]. These omics platforms function as comprehensive repositories for genomic, genetic, and related data resources concerning soybean. However, to date, no single database integrates data sets in a timely manner, which hinders the efficiency of data utilisation and application to soybean research. Additionally, omics analyzes frequently depend on specific reference genomes, which limits their applicability to other genomes. Providing easy-to-use tools to facilitate conversion between different genome coordinates is crucial. To address these issues, we developed a comprehensive multi-omics database structured on the available soybean datasets to provide an intuitive interface in a user-friendly, one-stop platform supported by online toolkits for functional gene mining in soybean.

The newly developed multi-omics database encompasses deep-sequencing data from 984 accessions, transcriptomic data across seed development in different varieties, extensive phenotypic data for genome-wide association study (GWAS) and phenotypic images from our laboratory. It integrates a wide range of omics datasets, including six modules “Genome, Phenome, Variome, Population, Transcriptome, and Synteny”, enabling rapid searching of genotypic and phenotypic data across germplasm or gene information. The database is named the SoyOD (Soybean Omics Database) and is available at https://bis.zju.edu.cn/soyod. The primary objective of developing this database and toolkit is to facilitate soybean functional studies and accelerate breeding efforts.

## Database content and features

### Overview of SoyOD interface

To establish a comprehensive repository of soybean multi-omics data, we meticulously integrated datasets from multiple sources to establish the soybean multi-omics database known as SoyOD. The SoyOD homepage provides an overview of features, module introduction, available tools, latest news and updates log. The top menu is a multi-omics search tool allowing users to input gene ID, chromosome region, genome name, germplasm name or phenotype of interest for the retrieval of information from the database. The detailed information of a gene can be found by viewing a related gene ID, including a basic description, gene structure and sequence, available quantitative trait locus (QTLs), and predicted functional categories. We have developed six interactive modules, namely “Genome”, “Phenome”, “Population”, “Transcriptome”, “Variome”, and “Synteny”, to integrate all datasets (**Figure** 1A). An analysis toolkit ensures interactive use between the different modules (Figure 1B). The homepage provides a drop-down menu and data summaries of the six modules, as well as links to “Toolkits”, “Download” and “About”. We have collected datasets from the last 20 years of global high-throughput sequencing projects, allowing all data to be accessed through a unified platform.

**Figure 1.**
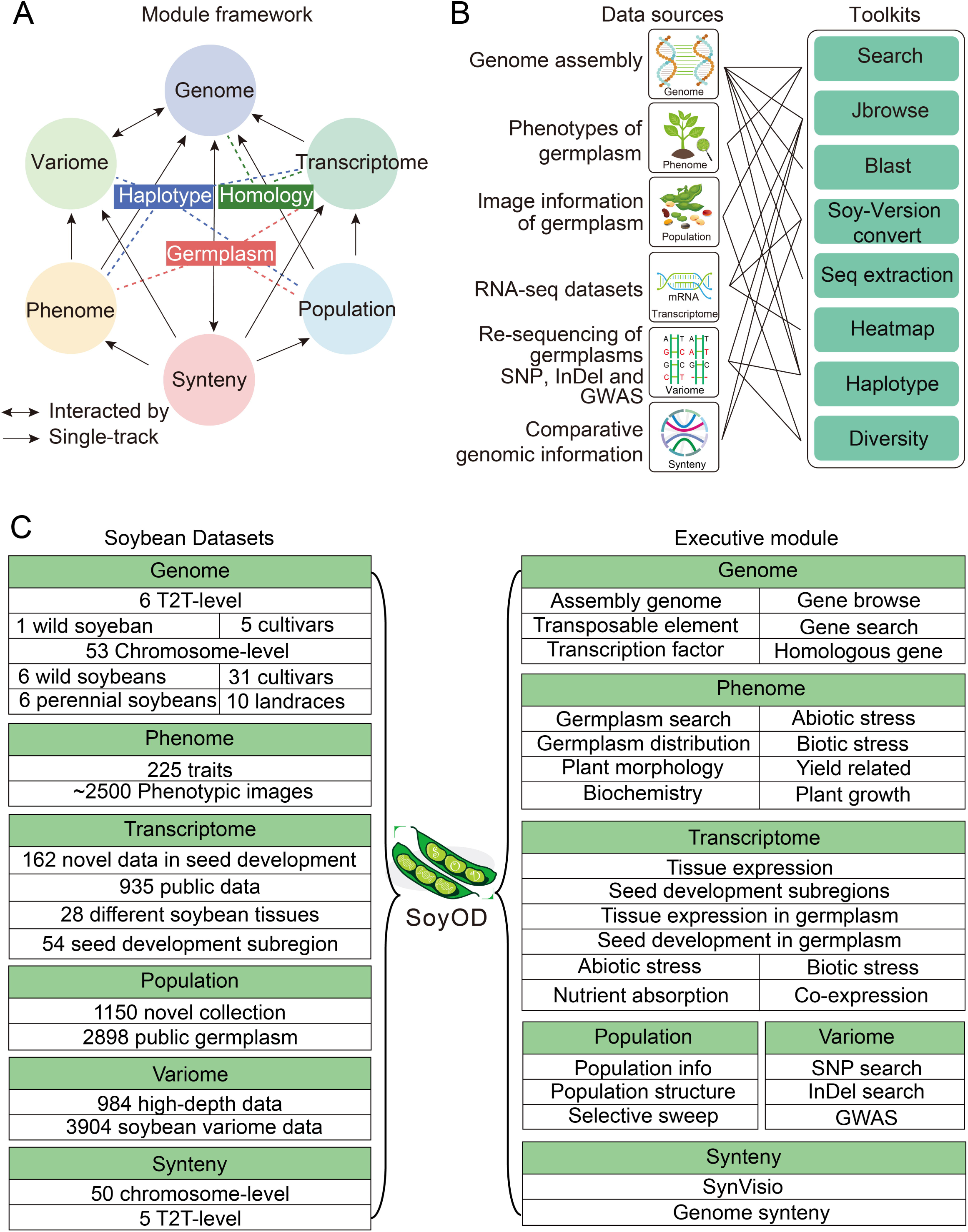
Overview of data sources, toolkits, and module interactions of SoyOD. **A.** Module framework of SoyOD, including genome, phenome, variome, population, transcriptome and synteny modules. **B.** The interactive interfaces between data sources and tools facilitate efficient data exchange and analysis. A connection line shows that this segment of the data can be linked or integrated with other parts. **C.** Description of soybean database datasets and introduction to the six executive modules and their submodules.

In the genome module, we collected 59 published soybean genome assemblies, which consisted of 6 perennial *Glycine* spp., 47 chromosome-level genomes (including 6 wild soybeans, 31 cultivars, 10 landraces) and 6 T2T-level genomes (including 1 wild soybean, 5 cultivars) (Figure 1; Table S1). The genome module includes several submodules, namely “Assembly genome”, “Gene browse”, “Gene search”, “Transcription factor”, “Transposable element” and “Homologous gene”, which users can easily search using gene ID or chromosome region (Figure 1C).

The phenome module contains 398,485 records of 225 phenotypes, including those related to plant morphology, yield related, plant growth, biochemistry, abiotic stress and biotic stress (Table S2). The phenome module contains phenotypic data for 4907 soybean germplasm resources and ∼2500 phenotypic images. These phenotypic data and images could be used for verification of a particular germplasm. Users can find the relevant phenotypic information and images according to a variety name or number search.

The population module combines the data for 940 newly sequenced accessions with 44 public datasets for a total of 984 deep-sequencing data with an average depth of 43×. In order to increase the size of the population scale, 2920 other genome sequences were integrated, for a total of 3904 population datasets. The high-depth re-sequencing set can be browsed or searched either separately or with the published re-sequencing sets. Furthermore, the selective sweep analysis serves as a foundation for users to identify genes associated with domestication and crop improvement.

Within the transcriptome module, a total of 1097 RNA-seq libraries were mapped to the corresponding genomes, and gene expression profiles were generated. Following classification, these datasets were organized into eight submodules, including Tissue expression, Seed development subregions, Tissue expression in germplasm, Seed development in germplasm, Abiotic stress, Biotic stress, Nutrient absorption and Co-expression (Figure 1C, Table S3). Five representative genomes are provided, including the two most recent T2T-level genomes (Williams 82 T2T and Zhonghuang 13 T2T) and three widely used chromosome-level genomes (Williams 82 v2, Williams 82 v4 and Zhonghuang 13 v2), are available as reference genomes. Users can conveniently retrieve gene expression data by entering the corresponding gene ID. All transcriptome raw data were downloaded and compared to multiple different assembled genomes to obtain read count, fragments per kilobase of exon model per million mapped fragments (FPKM), and transcripts per kilobase of exon model per million mapped reads (TPM) values. For more intuitive browsing of gene expression patterns, an electronic fluorescent pictograph (eFP) browser was developed for the visualization of gene expression levels from different dataset submodules. We have additionally created a heatmap toolkit that enables users to customize the expression matrix and generate personalized heatmaps. This functionality facilitates efficient access to gene expression data, ultimately enhancing the overall user experience. The co-expression network was constructed by the weighted correlation network analysis (WGCNA) package in R using datasets generated by different organizations and integrated into the co-expression submodule. Users can input one or more gene IDs into this submodule to investigate the co-expression network associated with specific genes of interest. Overall, the transcriptome module offers a comprehensive set of features and tools that enable users both explore level of gene expression in an intuitive interface.

We have gathered comprehensive variation data by resequencing a diverse of germplasm resources. The variome module includes a total of 984 high-depth sequencing datasets with an average depth of 43× (Table S4). All resequencing data were compared to the Zhonghuang 13 v2 reference genome. The high-depth sequencing of 984 accessions resulted in a total of 5,685,352 single nucleotide polymorphism (SNPs) and 1,361,946 insertions/deletions (InDels). In addition, the public resequencing datasets were analyzed and integrated, containing a total of 3904 accessions, comprising 7,193,573 SNPS and 753,361 InDels (Table S4). Users can query variation information within the input chromosome region. The variome module also includes GWAS analysis, which will be elaborated in greater detail below.

In the synteny module, we performed comparative genomic analysis of 56 assembled genomes (4 unannotated genomes were excluded). Users can readily find the structural variation (SV) information through comparison with the Zhonghuang 13 v2 reference genome. In addition, the platform incorporates a feature for quick browsing of genomic collinearity, utilizing the SynVisio web service (https://synvisio.github.io/). A user can select different chromosomes to browse by synteny and dot plot views. Meanwhile, users can quickly generate comparative genomic variation data by conducting searches across specified chromosome regions using the genome variation search modules.

### Germplasm resources and phenotypes

Germplasm resources are essential for crop improvement, biodiversity conservation, and research in plant genetics and breeding. These resources encompass a broad range of genetic material from plants, including seeds, tissue cultures, DNA, and other plant parts used for breeding new varieties or studying genetic traits. The SoyOD database includes phenotypic data of 4907 soybean germplasm resources and approximately 2500 phenotypic images. Users can search the germplasm data by species name, germplasm ID or trait name. The database provides detailed information for each germplasm, including phenotypic images and traits (**Figure** 2A) and measurements of important traits.

**Figure 2.**
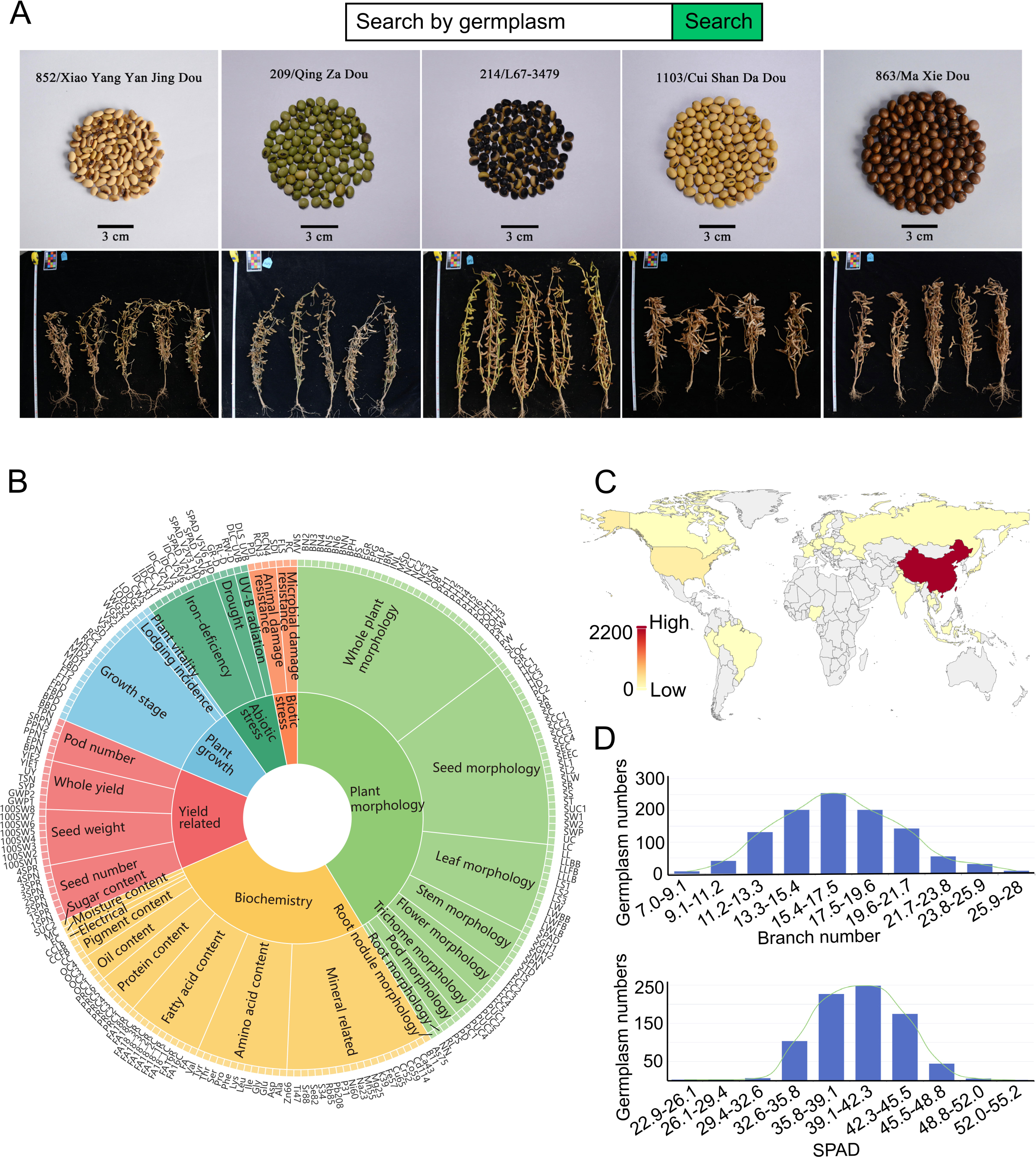
Phenotypic data retrieval module for soybean germplasm resources. **A.** Search bar and images gallery for soybean germplasm resources. **B.** Sunburst chart of 225 traits. **C.** Geographical distribution of germplasm resources in SoyOD. **D.** Distribution of branch number (upper) and SPAD value (lower).

A phenotype is an observable, physical characteristic of an organism. In the context of germplasm resources, phenotypic information is crucial for understanding how different genetic material reacts in various environments. This understanding informs breeding programs that are aimed at developing crops with desirable traits, such as high yield, disease resistance, and drought tolerance. In SoyOD, information pertaining to the source and phenotype was integrated into the phenome module. Phenotypic data were collected for 225 different traits, including newly collected data for 53 phenotypes and integrated data for 172 phenotypes from several other sources. The 225 phenotypes are classified into a three-level catalogue, and the user can select a phenotype by interacting with the sunburst graph (Figure 2B). The phenome module is divided into eight sub-modules (Germplasm search, Germplasm distribution, Plant morphology, Biochemistry, Plant growth, Yield related, Abiotic stress, and Biotic stress). The database contains a diverse array of germplasm resources from all over the world, including 3220 varieties from China (Figure 2C). The distributions of two traits, namely branch number (ranging from 7 to 28 branches) and soil and plant analyzer development (SPAD) values (ranging from 22 to 55) were plotted (Figure 2D). The germplasm interface facilitates rapid searches of accessions, allowing users to explore their characteristics with corresponding images, avoiding confusion between different varieties.

### Consolidation and interrogation of genomic data

In the genome module, we collected the 59 published soybean genomes, including 6 perennial *Glycine* spp., 47 chromosome-level genomes, and 6 T2T-level genomes (Table S1). The summary data on all 59 genome assemblies, including genome size, type, source, assembly level, and gene density, were integrated into the genome module. Complete annotation information (no gff file) was unavailable for 4 genomes and was consequently excluded from the partial analysis.

The mRNA, coding sequence (CDS), and protein sequence annotations are included in the gene browser, with each Gene ID linked to its gene structure, associated QTLs, and GO, KEGG, and PFAM predictions. Users can query and download these datasets by gene ID search. Within the submodules, users can browse transcription factors and transposons by assembly or by family name, while homologous genes can be browsed by Gene ID or by Homologous Group. The 73,270 groups were derived by comparing the predicted protein sequences from each genome. Users can query and download these datasets by gene ID search. Comparative genomic analysis between 56 genomes (excluding 4 without annotated information) and the Zhonghuang 13 v2 reference genome was conducted to generate synteny alignments and structural variation information, which are available in the synteny module.

### Population genetic variation detection and GWAS

The population and variome modules include 984 deep-sequencing datasets. In order to increase the population scale, another 2920 accessions were integrated, for a total of 3904 accessions. Comparison of the number of SNPs and InDels in the high-depth data and the 2898 published datasets [15]. Comparing the two datasets, multiple alleles and different alleles at the same variation site were removed, and a large number of shared SNPs (6382877) and shared InDels(631817) were found. Interestingly, there were 3,365,510 SNPs and 626,183 InDels that were only found in the 984 deep-sequencing datasets, suggesting that increased sequencing depth improves the detection of sequence variants (**Figure** 3B).

**Figure 3.**
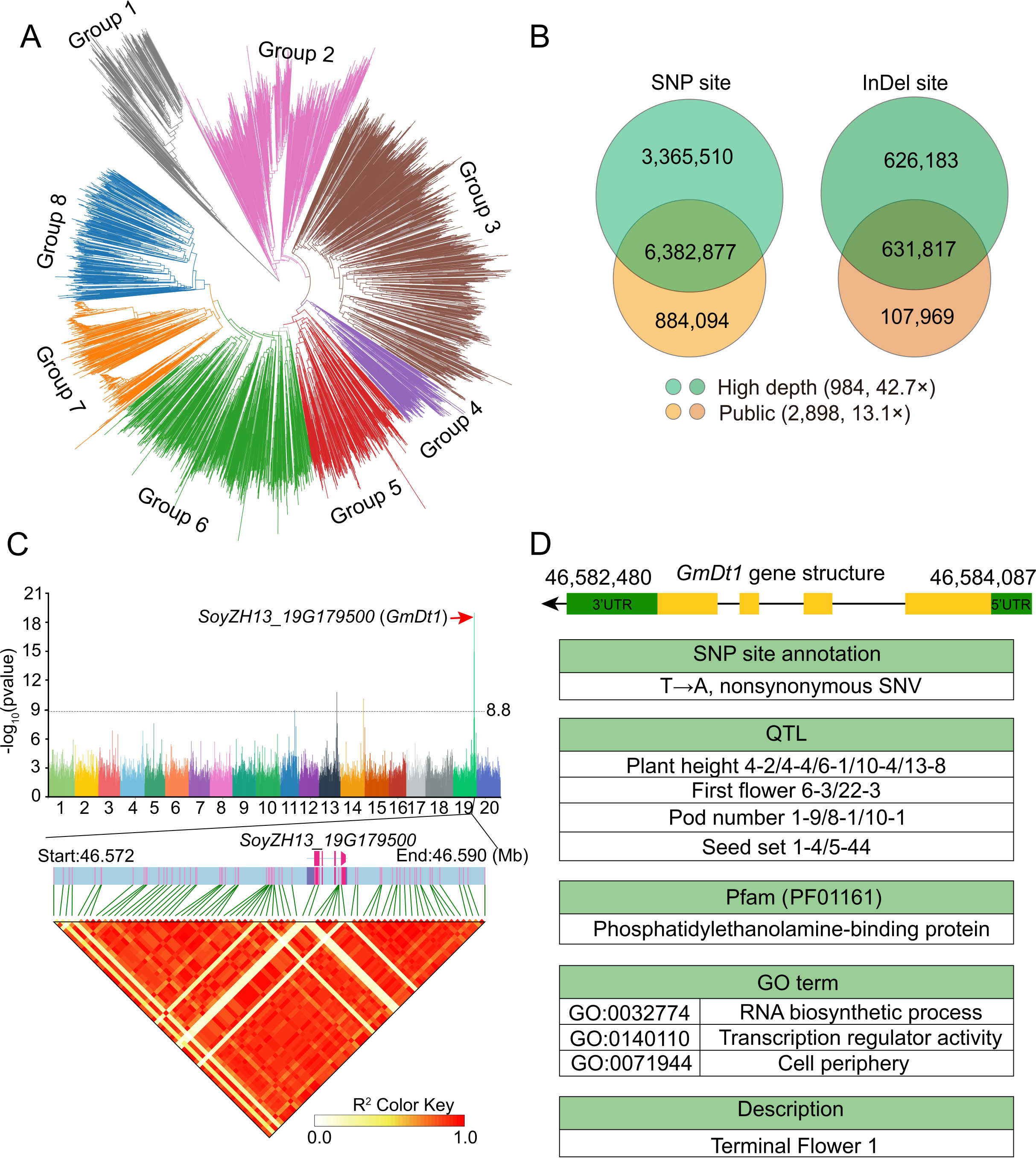
Gene mining via genome-wide association studies (GWAS) **A.** Phylogenetic tree of 3904 germplasm resources. **B.** The number of SNP and InDel sites detected by the high-depth sequencing (42.7×) from 984 accessions and 2898 public datasets (13.1×) using MAF < 1%, Miss > 10% as filtering criteria. **C.** GWAS of plant height using the efficient mixed model association (EMMAX) method as a case study, and analysis of the region by LDblock. **D.** Details of gene structure SNP annotation, linked QTLs, PFAM and GO terms of *GmDt1* retrieved from SoyOD.

GWAS analysis using the 984 resequencing datasets was conducted on 24 traits, including many related to plant morphology and yield. The GWAS submodule can be used to search for variations related to certain traits, we found a significant site (-log10(*p*) > 8.8) on chromosome 19 associated with plant height, which we hypothesized contained a dominant gene controlling plant height in soybean. This chromosome region contained a total of 849 loci. Through further screening, a significant locus in a predicted exon was found at Chr19:46582743, with an allele that leads to a non-synonymous substitution (Figure 3C). The linkage disequilibrium block (LDblock) under the haplotype tool also showed a strong linkage between this position and the preceding position. A search of the related gene in the main database search bar (which remains at the top of every SoyOD page), it was found that this site matches the gene *SoyZH13_19G179500* (*GmDt1*), which encodes a protein “Terminal Flower 1” (Figure 3D). By using the search function within the QTL section, the data can be filtered to highlight specific intervals linked to plant height, including the QTLs plant height 4-2, 4-4, 6-1, 10-4, and 13-8. *Dt1* was known to regulate stem growth habit and flowering in soybean [16]. The GWAS analysis in this study re-identified *GmDT1*, confirming the efficacy of the resequencing data and analysis methods for discovering novel genes.

### Gene functional exploration and visualized expression patterns

The SoyOD platform utilizes multi-omics approach to analyze population selection and visualize gene expression patterns, enhancing our understanding of complex biological phenomena. Here we present a comprehensive framework demonstrating how to perform soybean population selection analysis using an integrated method and visualize gene expression using an eFP or heatmap tools. *GmSWEET10a* has been reported to regulate soybean oil and protein content as well as seed size [17]. We use the SoyOD website to further illustrate the important role of *GmSWEET10a* in soybean seed development (**Figure** 4). For the case study, the population selective sweep submodule was used to view the data for Chr15 in the 3.85-4.05 Mb region. Four selective sweep models, namely the neutrality test statistics of fixation index (*F_ST_*), Tajima’s *D* statistic (Tajima’s *D*), cross-population composite likelihood ratio index (XP-CLR) and nucleotide polymorphism ratio (π ratio) values indicate that the gene region was under strong artificial selection during soybean domestication (Figure 4A). In addition, the QTLs cqSeed oil-007/010, seed protein 30-3, seed starch 1-3, seed volume 1-1, and seed length 1-1 were linked to this region (Figure 4B). These QTL descriptions are consistent with previous studies of the phenotypes [17]. In the coding region, four genetic variants were found, including two non-synonymous substitutions and one frameshift insertion, which may alter the sequence or structure of the SWEET protein (Figure 4B) and are therefore worthy of further investigation.

**Figure 4.**
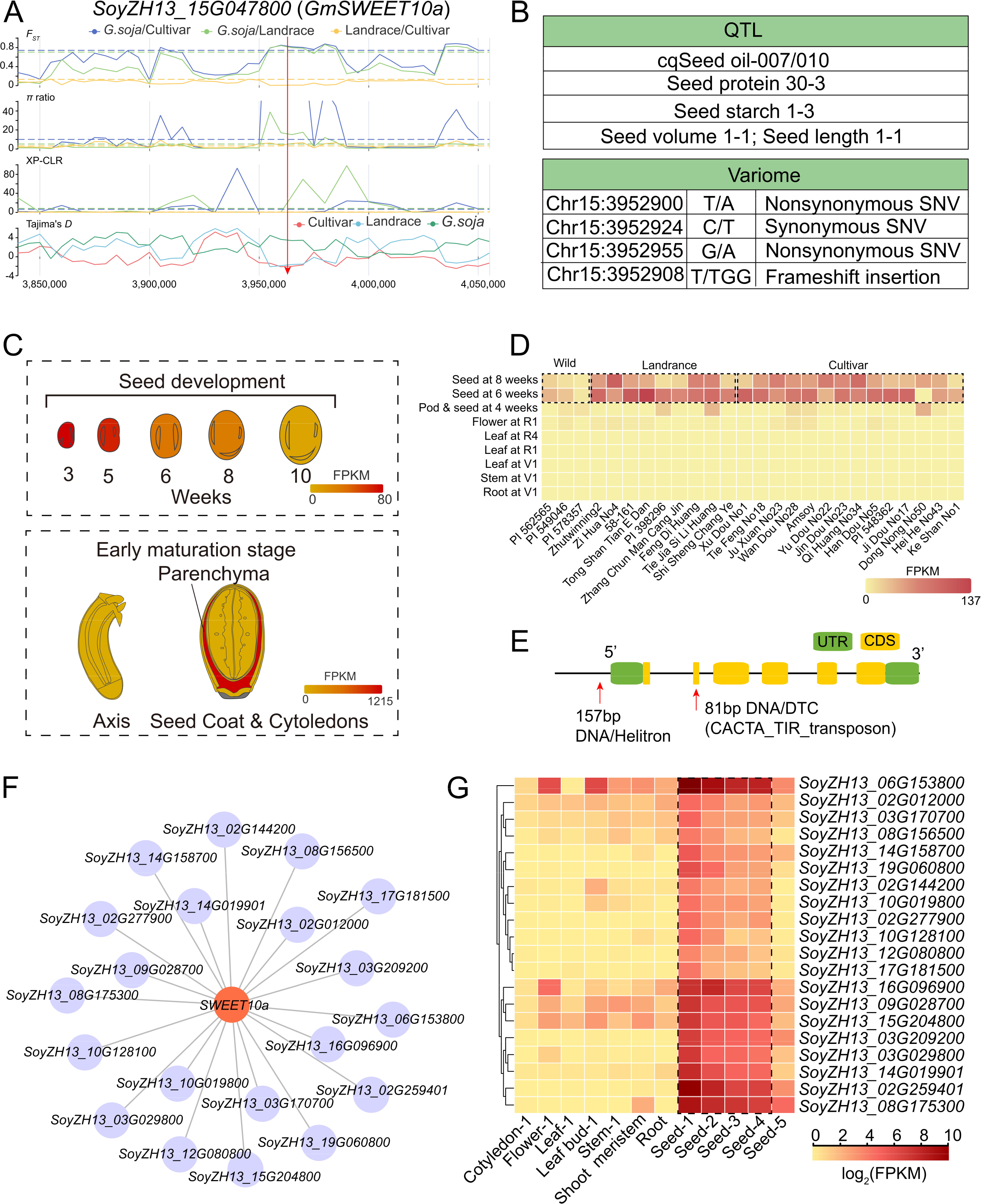
Case study characterizing *GmSWEET10a* using SoyOD. **A.** Genetic variation (*F_ST_*, π, XP-CLR and Tajima’s *D* values) was calculated between *G. soja*, landraces and the cultivars. **B.** *GmSWEET10a* QTL and variation information detected retrieved by the gene search submodule. **C.** Expression of the gene in the electronic fluorescent pictograph (eFP) viewer in the tissue expression and seed development submodules. **D.** Tissue expression in different germplasms. **E.** Transposon information for the *GmSWEET10a* gene, including both the gene body and the 2000 base pairs upstream of its transcription start site, using the transposable element module. **F.** Weighted gene co-expression network analysis (WGCNA) of *GmSWEET10a*. **G.** Expression profile of the co-expressed genes of *GmSWEET10a*.

The transcriptome data can be searched by selecting the Gene ID and corresponding genome. By searching the Zhonghuang 13 v2 genome with *SoyZH13_15G047800* (*SWEET10a*) and visualizing with eFP browser, the user can readily observe see a significant finding, high expression levels in early development stages (Figure 4C). Choosing the “seed development subregions” submodule, we found that the *SWEET10a* was predominantly expressed in the seed coat parenchyma at the early maturation stage (Figure 4C), underscoring its potential in shaping seed morphology and function. Moreover, when exploring the expression of *SWEET10a* in the ‘tissue expression in germplasm’ submodule, an intriguing pattern was evident in the heatmap. It was found that *SWEET10a* was specifically expressed in seeds, but at higher levels in landraces and cultivated accessions compared to the wild soybean accessions (Figure 4D). Such differential expression suggests that this gene may have been under selective pressure during domestication. Meanwhile, two DNA transposons within the genome region of *SWEET10a* were identified, located in the upstream and second exon regions of the gene, which may affect the function of the encoded protein (Figure 4E).

To identify genes that are co-expressed with *SWEET10a*, we performed co-expression analysis under the Transcriptome module. Twenty genes were associated with *SWEET10a* (Figure 4F). After downloading the list of co-expressed genes, we used the heatmap (under the Toolkits) to show the expression information of the 20 genes. These genes are selectively expressed during the early seed stage, with several showing high expression levels (Figure 4G). They are potential candidates for further functional analysis to explore gene interactions.

### Whole genome selective test and novel gene mining using SoyOD

Through the homology module, a user can find homologous genes within soybean genomes and *Arabidopsis*, as well as their functional descriptions, or use a BLAST search to find related information by target gene sequence (**Figure** 5A). A search using a gene ID will return target gene information including GO, KEGG, PFAM, QTL, sequence and gene structure data, tissue-specific, stress-responsive, or nutrient uptake-related expression profiles, and expression in different germplasm (Figure 5A). Once a user identifies a gene of interest, they can use the sequence to design a reverse genetics strategy to verify gene function.

**Figure 5.**
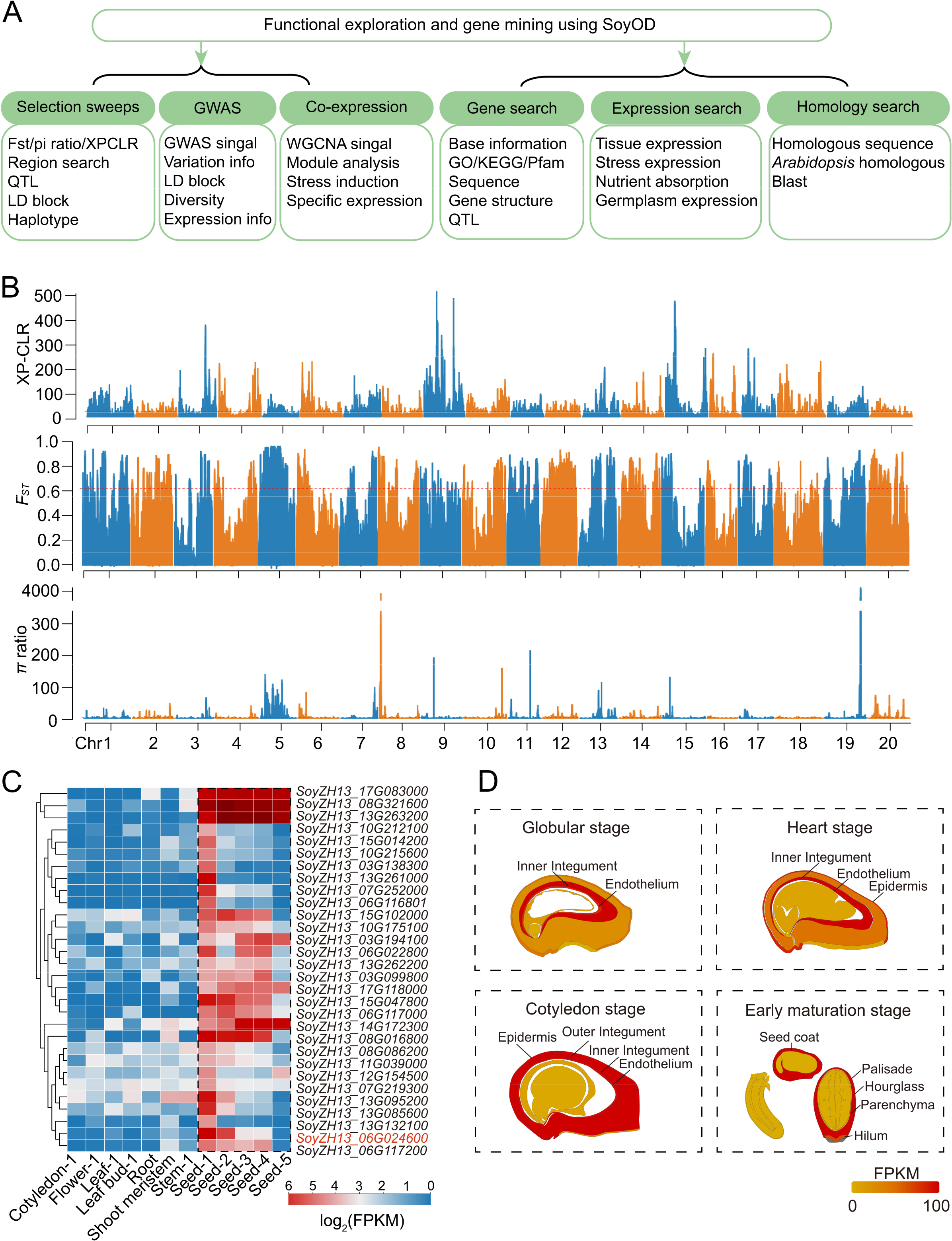
Discovery of new genes through SoyOD database exploration. **A.** Different functions of the SoyOD can be used for novel gene discovery.**B.** Selection signals analyzed using XP-CLR, *F_ST_*and π ratio, and to assess regions selected under domestication (from wild soybean to landrace). **C.** Regions under domestication selection overlapped with seed-specific transcripts. **D.** Expression of *SoyZH13_06G024600* in developing seeds. Data was extracted from Zhonghuang 13 v2.

Users can also access selective sweeps, GWAS, and co-expression data on SoyOD to search for genes altering a target trait (Figure 3, 5A). SoyOD contains a process for mining important genes based on population selection and expression data (Figure 5B). The population selection data relies on three measures, the *F_ST_*, π ratio and XP-CLR, and shows that soybean contains 1411 genes selected during domestication (Table S5) and 75 genes selected during improvement (Table S6), with 3 genes in both sweeps. Nine of these genes were found in regions previously reported as domestication regions. We combined these domestication regions with the tissue expression module to search for domestication genes expressed in specific regions of seeds. The 30 relevant genes were shown using the Heatmap toolkit (Figure 5C). We selected the example gene, *SoyZH13_06G024600*, for viewing in the seed development eFP module, and found that it was specifically expressed in inner integument of the seed coat (Figure 5D).

### Online omics interactive tool

We developed the toolkit portals on SoyOD to meet the growing demand for integrated multi-omics research. SoyOD hosts a diverse array of online analysis tools, including JBrowse, Blast, Soy-version convert, Seq extraction, Heatmap, Diversity and Haplotype (Figure 1). With access to sequences and genes from 59 (4 genomes were deleted without annotations) published soybean genomes in one place, the interactive tools on SoyOD significantly reduce the time researchers must spend navigating multiple databases. Users can conduct comprehensive analyzes and query various genomes through a single, unified platform, eliminating the requirement for switching between disparate databases. This innovation not only enhances efficiency but also fosters deeper insights by simplifying access to a broad array of genomic information. JBrowse allows users to easily query genomic annotation, variation and expression information, providing an overview of the data. A BLAST module, which can use an uploaded file, facilitates alignment of coding and protein sequences for comparisons between all of the annotated genomes. Users can select one or more genomes and enter multiple sequences simultaneously for convenient and rapid identification of homologous sequences. To convert between datasets, we have developed the Soy-version convert tool, which allows users to switch between different genomes. With the sequence extraction tool, a user can search for gene sequence information by gene ID or by region of the chromosome. Users can quickly and easily obtain expression information for multiple genes through the provided heat maps by entering multiple gene IDs and selecting the corresponding expression data, with the option to customize the matrix information output heatmap. Personalization of the heatmap can be used to adjust its maximum and minimum values and colors. The population diversity and haplotype tools allow users to better explore whether a gene is strongly selected.

## Discussion

Cultivating new soybean varieties to enhance yield and quality is crucial for meeting the demands of the soybean market. Functional genomics and molecular breeding in soybean increasingly rely on multi-omics analysis based on existing datasets [18–20]. To support genetic improvement and molecular breeding of soybean, we integrated six modules, the Genome, Phenome, Variome, Population, Transcriptome, and Synteny into an easily usable, comprehensive multi-omics database named SoyOD. During this process, we included and compared the multiple omics datasets generated over the past decade (Figure 1; **Table 1**). Compared with the available databases, SoyOD has several advantages: 1) SoyOD contains the largest and most up-to-date soybean omics datasets, including 59 assembly genome datasets, 398,485 records of 225 phenotypes, 1097 RNA-seq transcriptomes, genetic variation data from 3904 soybean accessions, selective sweep information, and 59 genome synteny and comparative genomic analysis. This makes SoyOD the most comprehensive resequencing soybean population ever studied. 2) SoyOD offers new high-depth resequencing data from this study that can be used for more precise genetic variation analysis. 3) SoyOD contains phenomic data of soybean germplasms, which is crucial for understanding genotype-phenotype relationships. 4) SoyOD implements multi-module interactive use, making the interface user-friendly and more intuitive. Users can input a gene ID to search the corresponding genome location, gene annotations and predicted translation and related QTL information. The comparison of SoyOD with other available soybean genome database is listed in Table 1.

**Table1.** Comparison of current popular soybean databases.

| Website | Assenbly genome | Sequenced No. accession | RNA-seq libraries | Traits | Phenotype images | Epigenetics | GWAS | Variation phenome | Novel data |
| --- | --- | --- | --- | --- | --- | --- | --- | --- | --- |
| SoyOD | 59* | 3904 | 1097 | 225 | ~2500 | - | √ | √ | √ |
| Soybase | 14 | # | - | - | - | 6 | - | - | - |
| SoyKB | 3 | 775 | - | - | - | 4 | √ | - | - |
| SoyFGB | - | 2214 | - | 42 | - | - | √ | - | - |
| SoyOmics | 29 | 2898 | 81 | 115 | 28 | 81 | √ | √ | - |
| SoybeanGDB | 39 | 3379 | 146 | - | - | - | - | - | - |
| SoyGVD | - | 4414 | - | 48 | - | - | - | - | - |
| SoyMD | 38 | 4414# | 333 | 120 | - | 242 | √ | √ | - |

Note: \*indicates 6 T2T-level genomes; # indicates including SNP50k data; - indicates that the data is not included; √ indicates that the data is included.

With the accelerated development and application of sequencing technology in the post-genomic era, gene mining and molecular breeding increasingly rely on multi-omics analysis [20]. SoyOD integrates the majority of multi-omics research data into a user-friendly interface with a variety of online analysis tools. SoyOD is designed as a dynamic platform to integrate the latest global soybean multi-omics data. Notably, SoyOD is accessible online without requirement for registration, making it readily available to researchers. While we recommend Chrome browser for the best browsing experience, the database is optimized to function across various browsers. In the future, we will continue to update this database and develop analytical methods and tools to make it an important resource for global soybean research.

## Materials and methods

### Data acquisition

To construct an accessible, comprehensive multi-omics database for soybean, we developed six interactive modules including “Genomic”, “Phenomic”, “Population”, “Transcriptomic”, “Variomic” and “Synteny”. In the genome module, we collected the data from 59 previously reported, high-quality assembled genomes, including 47 chromosome-level genomes [6 wild soybean (*Glycine soja*), 10 landraces, 31 improved cultivars], 6 T2T-level assemblies of the genomes from Zhonghuang 13 (including two versions namely T2T and T2T-2), Williams 82, , Yundou1 and Jack, and 6 recently published genomes from different perennial soybeans species (*Glycine cyrtoloba*, *G. dolichocarpa*, *G. falcata*, *G. stenophita*, *G. syndetika*, and *G. tomentella D3*) [2, 5–9, 15, 21–27]. The publication for each genome is listed in Table S1. However, 4 genomes (Zhonghuang 13 T2T-2, ENREI, PI594527 and Zhe Nong 6) did not contain annotation information, and only partial genomic analyses were performed.

The phenome module collects 398,485 records of 225 phenotypes obtained from 53 local data resources and 172 public data resources over multiple years. All traits were divided into six different phenotypic classes, namely plant morphology, yield-related, plant growth, biochemistry, abiotic stress and biotic stress.

Transcriptomic data were collected from 1097 RNA-seq libraries from different tissues and conditions, including seed development stage, abiotic stress, biotic stress and so on (Table S3).

To determine the expression patterns of various soybean varieties throughout seed development, we sequenced 162 RNA-seq libraries. These libraries were derived from 54 different accessions, which were sampled at four key stages of seed development from 15 representative soybean germplasm resources. In addition, we downloaded and integrated 342 expression datasets from 28 tissues from cultivar Williams 82 [28], 27 tissues from cultivar Zhonghuang 13 [4] and 9 tissues from 26 other varieties [15]. We included data from 297 accessions of soybean seeds sampled from different stages of early seed development by laser capture microdissection [29, 30]. In the abiotic stress submodule, 296 libraries were taken from plants subjected to different conditions, including exposure to salinity, drought, cold, temperature, flooding, dehydration, PEG 8000, submergence, pH, CO_2_, and auxin treatments. In the nutrient absorption submodule, treatments included nitrogen, phosphorous, potassium, iron and zinc. References for these datasets are listed in Table S3.

In variome and population modules, we combined 940 newly sequenced, high-depth soybean genomes and 44 published high-depth resequencing genomes (with an average depth of 43×) to construct high-quality genome maps based on second-generation sequencing and phenotypic data of representative germplasms. With the addition of 2920 public resequencing datasets [15, 23, 24], the total resequencing data included 3904 datasets (Table S4).

### Gene annotation and function analysis

The six T2T and 53 chromosome-level genomes listed in Table S1 were integrated. The 56 (4 unannotated genomes were excluded) genomes were annotated using eggNOG with default program settings, and the GO, KEGG functional descriptions and PFAM domains [31]. Prediction of transcription factors (TF) used protein homology alignment to *Arabidopsis* (https://www.arabidopsis.org/) or soybean transcription factors by the Plant TFDB [32]. Transposons of 59 genomes were predicted using the EDTA pipeline [33] by detecting repeat sequences with the default parameters.

### Variation calling and annotation

The sequenced 984 high-depth genomes were compared to the Zhonghuang 13 v2 reference genome [3] using the Burrows-Wheeler Aligner (BWA; v 0.7.17-r1188) [34]. SAMtools (v1.14) [35] and Picard package (http://broadinstitute.github.io/picard/, v2.27.5) were used for duplicated reads. The detection was performed using the GATK (v4.2.2.0) HaplotypeCaller [36], genotyping was done with GenotypeGVCFs [36], and the separation of SNPs and InDels was performed using the GATK SelectVariants option [36]. Finally, low quality data was filtered for SNP and InDel annotation using ANNOVA software and removed [37]. An additional 2898 germplasm resources were downloaded from the National Genomics Data Center (NGDC) [15], For subsequent analysis, all data was filtered and annotated. The public re-sequencing data was analyzed according to the same method, so that the module contains variation data from a total of 3904 accessions.

### Selective sweep and GWAS analysis

To analyze the population structure and construct an evolutionary tree, we screened a subset of SNPs and InDels using VCFtools (v.0.1.16) [38]. The cutoff settings for the data from the 3904 re-sequenced accessions were a missing rate of > 10% and a minor allele frequency (MAF) of < 1%; for the 984 accessions, the SNP missing rate was > 10% and the MAF was < 5%, while the InDel missing rate was > 50% and the MAF was < 1%. The evolutionary trees were constructed using Phylip software (https://phylipweb.github.io/phylip/index.html) with 1,000 bootstraps. Population structure was analyzed using the admixture (v1.3.0) program with the parameters k = 2 to k = 10 according to phylogenetic analysis order. To analyze each subpopulation, we employed VCFtools and XP-CLR (updated based on Python) [39] to calculate the genetic diversity (π), Tajimas’ *D*, XP-CLR and *F_ST_* values using a window size of 20 kb and a step size of 2 kb across the entire soybean genome. The top 5% value was chosen as the significance threshold.

GWAS were performed using the EMMAX algorithm [40] using SNP data. For GWAS, the threshold was set at 0.05/(total number of SNPs), corresponding to -log10(*p*) = 8.1 and 0.01/(total number of SNPs), corresponding to -log10(*p*) = 8.8 as whole-genome significance cutoff. Manhattan and QQ plots were drawn by CMplot (R package). LDBlockShow was used to draw LD block heatmaps for a specified area using the default parameters [41].

### Comparative genome analysis

Genome sequences of Zhonghuang 13 v2 were compared with those of 55 other assemblies (excluding 4 unannotated ones) using the NUCmer program (version 4.0.0rc1) within the MUMmer4 suite [42]. Following the filtering one-to-one alignments to retain only those with a minimum length of 100 bp (utilizing the delta-filter program from MUMmer with the specified parameters ‘-m -i 90 -l 100’), the show-coords program was used to convert the delta file into readable matching coordinates. Syri (v1.6.3) [43] was employed to extract the coordinates and features of structural variations. Finally, the plostr program from Syri was used to generate visual results. Gene synteny was identified through the use of the Python-based McScan tool[44]. For visualization of the McScan results, the dotplots and synteny were drawn in SynVisio [45] before integration into SoyOD.

A total of about 494,137 genes from the 56 soybean genome assemblies utilized to construct gene clusters. All protein sequences were obtained and homologies were detected using the OrthoFinder (v2.5.5) software (with the -S diamond -M msa -T fasttree parameters). Homologous genes were aggregated into clusters, yielding a total of 73,270 groups.

### Transcriptomic analysis

A total of 1097 transcriptome libraries were downloaded using the prefetch (v2.8.0) software, and then adapter sequences and low-quality information were removed using fastp (v0.12.4) [46] (Table S3). To obtain a comprehensive gene expression profile, we compared datasets to 5 frequently-used reference genomes (Zhonghuang 13 v2, Zhonghuang 13 T2T, Williams 82 v2, Williams 82 v4 and Williams T2T), while other RNA-seq datasets were compared to the genome corresponding background (Table S1) by Hisat2 (v2.2.1) software with default parameters [47]. The gene expression levels were quantified to obtain count, FPKM, and TPM values using the ‘featurecount’ R package [48]. The FPKM values were then analyzed by WGCNA [49] for co-expression analysis using topological overlap matrix (TOM) program. Any expression value less than 0.25 was excluded. The resulting co-expression network was obtained through the calculation of the Pearson correlation coefficient between pairwise gene expression levels.

### Website construction and design

SoyOD (https://bis.zju.edu.cn/soyod) is a web-based application built on the Python Django (v2.2.14) framework for the backend and HTML, CSS and jQurey (v3.4.1) as the frontend JavaScript framework, ensuring a powerful and flexible foundation for data processing and management. Data storage and retrieval are managed through an Apache 2 web server (v2.6.1) and uses MySQL (v8.0.35) as its database engine. The visualization software included Echarts (v5.4) [50] for line, bar, pie charts, etc; D3 (v6.7.0) [51] for evolutionary trees; JBrowse2 (v2.8.0) [52] as a genome browser; and Diamond (v2.0.14) for gene sequence BLAST analysis. The whole system uses the Ubuntu operating system (v20.04; Canoncial Ltd.). Some of the icons were designed using Adobe Photoshop (v19.0) and Adobe Illustrator (v27.0). The database is accessible online without registration and is recommended in Google Chrome, Microsoft Edge, Mozilla Firefox or Apple Safari. SoyOD uses this combination of technologies to create an intuitive and user-friendly interface and to deliver a seamless experience for accessing soybean-related data and tools.

## Data availability

The raw RNA-seq data used in this study were deposited in the National Genomics Data Center (https://ngdc.cncb.ac.cn/) Genome Sequence Archive (GSA) database, registration number for PRJCA030195. SoyOD is publicly available at https://bis.zju.edu.cn/soyod.

## CRediT author statement

**Huixia Shou**: Conceptualization, Data curation Funding acquisition, Resources, Project administration, Supervision, Writing - review & editing. **Jie Li**: Conceptualization, Data curation, Formal analysis, Investigation, Methodology, Software, Validation, Visualization, Writing - original draft, Writing - review & editing. **Ming Chen:** Conceptualization, Data curation, Project administration, Resources, Supervision, Writing - review & editing. **Qingyang Ni**: Data curation, Formal analysis, Validation, Visualization, Methodology. **James Whelan**: Funding acquisition, Project administration, Writing - review & editing. **Guangqi He**: Data curation, Validation. **Jiale Huang**: Validation, Data curation. **Haoyu Chao**: Methodology. **Sida Li**: Data curation. **Guoyu Hu**: Resources, Investigation.

## Acknowledgments

We thank and Dr. Jiming Xu, Shelong Zhang (Zhejiang university) for technical assistance, and Zhenyu Qi and Rui Sun of Agricultural Experiment Station of Zhejiang university for plant propagation and field test. The authors also thank Ms. Anita K. Snyder for her assistance with language checking. This work was supported by the Ministry of Science and Technology of China (2021YFF100204), HS; the Zhejiang Provincical Key Research and Development Project (2021C02057); The Zhejiang Provincial government Kun Peng Fellowship. All authors have reviewed and consented to the final version of the manuscript.

## Supplementary material

**Table S1** Chromosome- and T2T-level genome assemblies collected and used in SoyOD.

**Table S2** Summary of phenotypic datasets.

**Table S3** Summary of collected transcriptome datasets.

**Table S4** The 3904 re-sequenced accessions of *Glycine soja* and *Glycine max*.

**Table S5** Screening of selective signals for domestication-related genes.

**Table S6** Screening of selective signals for improvement-related genes.

